# RECoN: Rice Environment Coexpression Network for Systems Level Analysis of Abiotic-Stress Response

**DOI:** 10.1101/166694

**Authors:** Arjun Krishnan, Chirag Gupta, Madana M.R. Ambavaram, Andy Pereira

## Abstract

Transcriptional profiling is a prevalent and powerful approach for capturing the response of crop plants to environmental stresses, e.g. response of rice to drought. However, functionally interpreting the resulting genome-wide gene expression changes is severely hampered by the large gaps in our genomic knowledge about which genes work together in cellular pathways/processes in rice. Here, we present a new web resource – RECoN – that relies on a network-based approach to go beyond currently limited annotations in delineating functional and regulatory perturbations in new rice stress transcriptome datasets generated by a researcher. To build RECoN, we first enumerated 1,744 stress-specific gene modules covering 28,421 rice genes (>72% of the genes in the genome). Each module contains a group of genes tightly coexpressed across a large number of environmental conditions and, thus, is likely to be functionally coherent. When a user provides a new differential expression profile, RECoN identifies modules substantially perturbed in their experiment and further suggests deregulated functional and regulatory mechanisms based on the enrichment of current annotations within the predefined modules. We demonstrate the utility of this resource by analyzing new drought transcriptomes of rice in three developmental stages, which revealed large-scale insights into the cellular processes and regulatory mechanisms involved in common and stage-specific drought responses. RECoN enables biologists to functionally explore new data from all abiotic stresses on a genome-scale and to uncover gene candidates, including those that are currently functionally uncharacterized, for engineering stress tolerance.

## Introduction

The complex response of plants to abiotic-stress spans several orders of magnitude in time and space, causing system-wide adverse reactions and protective responses. Gene expression profiling has been used successfully to capture the system-wide molecular programs that underlie the cellular response to abiotic stresses (Deyholos, 2010). Analyses of the drought-stress inducible transcriptome in Arabidopsis, for example, reveal a plethora of responses including the induction of transcription factors, phospholipases C and D, protein kinases (MAPK, CDPK), proteinases, water channel proteins, antioxidant enzymes and molecules (GSTs, thioredoxins, peroxiredoxins), factors such as chaperones that afford protection for macromolecules (LEA proteins, HSPs) and osmoprotectant synthases (for proline, betaine, sugar) (Seki et al., 2002; Shinozaki et al., 2003; Harb et al., 2010). Making such analytical inferences from the transcriptome hinges on the availability of prior functional and regulatory knowledge about a large number of genes in the genome, which can then be used to meaningfully summarize genome-wide gene-expression changes. Although far from complete in Arabidopsis, such functional/regulatory information about genes – what they do, how they work together, and how they are regulated – is severely lacking in rice both in quality and genomic coverage. This paucity has led to a significant lag in the number of characterized drought-responsive genes and cellular processes in rice compared to Arabidopsis. Therefore, we need alternative approaches that can go beyond currently available gene annotations for fully extracting knowledge from rice transcriptomes and help towards gaining a comprehensive understanding of stress-response in this crop plant.

A promising starting point for such an alternative approach is the large amount of currently publicly available gene expression data in plants. A powerful analysis framework that has emerged in recent years involves estimating the similarity of expression patterns between all pairs of genes across diverse conditions to build gene coexpression networks, representing the genome-wide transcriptional organization of the cell (Usadel et al., 2009; Pearce et al., 2015). Particularly, in the crop model rice, recent studies have used gene coexpression to gain biological insights into general (Wang et al., 2009; Shaik and Ramakrishna, 2013) and case-specific (Fu and Xue, 2010; Ambavaram et al., 2014) gene regulation. Coexpression networks have also been used extensively in plants to organize genes into transcriptional modules and explore their functions (Mentzen and Wurtele, 2008; Mao et al., 2009). These modules represent groups of genes/proteins are likely to work together to perform a coherent biological function inside the cell (Hartwell et al., 1999), essentially expanding upon the available functional annotations. Hence, coexpressed modules can be used as sets of functionally coherent genes to see their enrichment in new expression data, especially in genomes where the functional annotations are sparse and incomplete. However, we still lack such a resource that allows enrichment analysis of coexpressed clusters/modules in new expression data of rice.

Therefore, it would be valuable to reconstruct a rice coexpression network that integrates information across a large number of datasets specifically in the context of stress response. In addition, it would be highly beneficial if researchers could bring this coexpression network to bear on their new gene-expression profiles (in one or more stress conditions) for functional resolution and comparison. Interpreting the long lists of responsive genes, a typical result of a gene-expression study, will become amply tractable by identifying subsets of responsive genes that are likely to be functionally coherent. Likewise, comparison between gene responses in different growth stages or conditions is likely to be more meaningful and robust at the level of cellular functions/pathways than at the level of individual genes (capturing the perturbation of different subsets of the same cellular apparatus and overcoming the various sources of noise in high-throughput assays).

To meet all these critical needs, here we present a genomic resource for comprehensive analysis of stress response in rice based on a modular coexpression network specific to response to environmental conditions, and apply this resource to perform a detailed analysis of stage-specific drought response in rice. First, we carried out a genome-scale analysis integrating publicly available rice gene expression datasets generated in the context of response to a range of environmental conditions. Next, using this integrated data, we constructed, what is termed, the Rice Environment Coexpression Network (RECoN), based on gene expression correlation across environmental conditions. Finally, we partitioned RECoN into densely connected modules using a graph-clustering algorithm. As a pertinent test case for our approach, we performed gene expression profiling of rice plants subjected to drought at three developmental stages. We used this data to perform both a traditional analysis – functional analysis (using Gene Ontology) – and a new analysis using RECoN, teasing out drought-related modules within the drought-response genes identified from our experiments. The new RECoN-based analysis of new experimental data helped highlight pathways, processes, regulatory genes, and potential transcriptional regulatory mechanisms critical for drought response in rice. We have made RECoN available for rice stress biologists through an interactive network browser at https://plantstress-pereira.uark.edu/RECoN/. Biologists can use this resource to explore coexpression clusters within their stress transcriptome and systematically guide follow-up experimental studies for constructing the underlying gene network.

## Materials and Methods

### Coexpression Network Analysis

A total of 29 publicly available gene expression datasets comprising of 414 samples of the Affymetrix rice GeneChip from were collected from NCBI GEO (Barrett et al., 2009) and ArrayExpress (Parkinson et al., 2009). From these, 129 samples (45 groups) with a unifying biological theme, i.e. response to some environmental condition, were used for coexpression analysis (see Supplementary Table S1).

We previously reported the re-annotation of Rice GeneChip to increase the reliability of expression quantification (Ambavaram et al., 2011). Briefly, the chip definition file (CDF GPL11322) was created by mapping probes to target genes that have perfect sequence similarity, and regrouping probesets such that each represents a single corresponding gene. We used this custom CDF to background correct, normalize and summarize the raw data using justRMA (Irizarry et al., 2003), with values in replicate samples averaged. To estimate coexpression, the Pearson correlations between every pair of genes were first calculated (Huttenhower et al., 2008) and then normalized using Fisher’s Z-transform (David, 1949). Then, the standardization of these scores resulted in coexpression score (*z_cs_*) indicating the number of standard deviations it lies from the mean, and follows a normal distribution to be interpretable by the level of significance, as |*z_cs_*| values greater than 1.96 allowed a 95% confidence interval to work with.

A coexpression network was then constructed connecting pairs of genes that have a *z_cs_* >1.96 (top 2.5% of all pairs of genes ordered in decreasing order of correlation). This cutoff corresponded to a Pearson correlation coefficient of 0.632. This network that contains 34,792 genes connected by ~18.5 million edges was then clustered using SPICi (Jiang and Singh, 2010). Since SPICi requires a density parameter *T_d_* as input, a range of values of the parameter from 0.1 to 0.9 was tested. Clusters obtained using each *T_d_* value were evaluated using several criteria including the number of clusters formed, fraction of genes in clusters of size 3 or more, average segregation (modularity), and extent of overlap between clusters and GO BP gene sets (termed ‘functions’) (Fig. 6). In order to calculate average segregation, as desired property of dense interaction networks, the coexpression network is modeled as an undirected graph *G=(V, E)*, consisting of a set *V* of nodes (i.e., genes) and a set *E* of edges (i.e., coexpressing gene pairs). Let *w_uv_* denote the weight of the edge (*u, v*) ∈ *E*, denoting the Pearson correlation coefficient of gene pairs (*u, v*). The graph *G_c_=(V_c_, E_c_)* is defined as the graph induced by the genes that are part of cluster *c*, and average segregation is computed as:

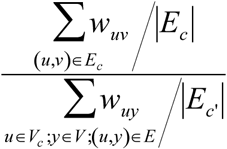

where *E_c’_* is the set of edges in *G* that are incident on *V_c_*. For functional enrichment analysis the overlap between genes within a cluster and genes annotated to a given GO BP term using the cumulative hypergeometric test. Using only GO BP terms that annotate <500 genes (to ensure a certain level of specificity in definition), for a pair of gene sets (cluster and GO BP term) *i* and *j*, if *N* is the total number of genes, *n_i_* and *n_j_* are the number of genes in gene set *i* and *j*, and *m* is the number of genes common to the gene sets, the probability (*p*-value) of an overlap (enrichment) of size equal to or greater than observed is given by the formula below.

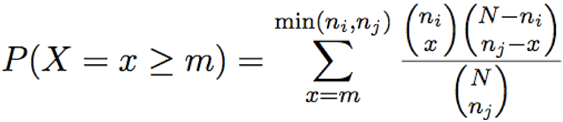

*P*-values from the test were converted to *q*-values to correct for multiple hypothesis testing using Benjamini-Hochberg method (Benjamini and Hochberg, 1995) and cluster-GO_BP pairs with *q*-value <0.1 were considered for analysis. The level of functional enrichment in a cluster is quantified using –*log_10_*(*q*-value).

After clustering the network using SPICi with *T_d_* value of 0.65, from all the clusters, those relevant to drought were determined by testing which clusters contained a significantly high number of drought-regulated genes up- or down-regulated in any one of the stages (again using a cumulative hypergeometric test). Then, four sets of genes were extracted from each ‘drought’ cluster – all the genes in the cluster, and seedling, vegetative and reproductive drought-regulated genes – for discovery of putative cis-regulatory elements and enrichment analysis of GO biological processes in the rice genome.

### Rice Plant Material and Drought-Stress Treatments

Rice (*Oryza sativa* L. ssp *Japonica* cv. Nipponbare) seeds were germinated in hydroponic half-strength Hoagland solution and seedlings were grown about a week in an environmentally controlled growth chambers maintained at 28^0^C ± 1 temperature, 65% relative humidity with a daily photoperiodic cycle of 14h light and 10h dark, and then plants allowed to reach the reproductive stages were grown in soil under greenhouse conditions. Samples of well-watered and drought stressed were collected at various developmental stages which include seven-day old seedlings, vegetative (V4) and reproductive (R4) stages based on discrete morphological criteria as described by Counce et al (2000).

For drought treatment, plants were gradually subjected to field drought stress in order to reach 50% field capacity (FC) by regulating the water supply, whereas control plants were maintained at 100% FC. During the stress period the pots were weighed daily and the difference in weight on subsequent days was corrected by adding water to maintain the required field capacity. The physiological condition of plants at 50% FC was monitored by chlorophyll fluorescence, quantum yield (Fv/Fm) and the relative water content (RWC) (Supplemental table S1). For dry down drought treatment, rice plants after transplanting were separated, with 5 pots maintained at well-watered condition serving as control while another set of 5 pots were used for drought experiments. For drought stress treatment water was withheld until the moisture level progressively dropped down to 6%. Drought stress symptoms were monitored for leaf rolling and measurement of soil moisture content everyday using soil a moisture meter (Rapitest). For all the stages (seedlings, vegetative and reproductive), three biological replicates were harvested from independent populations of plants, when leaves were completely rolled and RWC was around 65-70%. RWC was measured in the leaves used from where photosynthesis was measured. Leaf fragments of same size were cut and fresh weight was measured and hydrated immediately to full turgidity in deionized water for 6 h. After 6 h the leaf fragments were blotted on paper towels and the fully turgid weight was taken. Turgid leaf samples were then oven dried at 80°C for 72 h and weighed to determine dry weight. RWC percentage was measured as: RWC (%) = (fresh weight − dry weight)/ (turgid weight − dry weight) × 100. The drought stress symptoms such as leaf rolling and basal leaf senescence were apparent in stress-induced plants, while control plants growing at 100% FC were observed to grow well showing 95% RWC.

### Measurement of Chlorophyll Fluorescence and Quantum Yield

Chlorophyll fluorescence and the quantum yield was measured by using the Modulated Chlorophyll Fluorometer OS1-FL (Opti-Sciences Inc, USA). During and after stress treatments, flag leaf from stressed and unstressed wild-type was placed in close contact with the Photosynthetically Active Radiation (PAR) clip, which provides basic data to the OS1-FL system on ambient conditions. The PAR sensor is designed to measure leaf temperature and the light intensity. The ratio of variable fluorescence (Fv/Fm) and the yield of quantum efficiency (Y) are indicative of photosystem I and II performance of the plants under stress.

### RNA Isolation, Probe Labeling, and Hybridization

Total RNA was isolated from the rice seedlings, vegetative (V4) and reproductive (R4) tissues of both control and stress treated plants using the RNeasy plant kit (Qiagen, USA) according to manufactures protocol. For each stage/treatment, three independent biological replicates were used for RNA isolation.

RNA quantity, quality, and purity were assessed with the use of the RNA 6000 Nano assay on the Agilent 2100 Bioanalyzer (Agilent Technolgies, USA). Total RNA (~4 µg) from each sample was used to generate first-strand cDNA with a T7-Oligo(dT) primer. Following second-strand synthesis, in vitro transcription was performed using the GeneChip^®^ IVT Labeling Kit according to the manufacturer’s instructions. The preparation and processing of labeled and fragmented cRNA targets, as well as hybridization to arrays, washing, staining, and scanning were carried out according to manufacturer’s instructions (http://www.affymetrix.com). The Affymetrix Rice GeneChips (which contain~ 43,000 probe sets or genes), washing and scanning were carried out in Gene chip fluidics Station 450 (Affymetrix) and the Gene chip Scanner 3000 by Affymetrix (Santa Clara, CA, USA), respectively.

### Analysis of Differential Gene Expression

The custom CDF file was used to background correct, normalize and summarize all the raw expression data using RMA in R (Ihaka and Gentleman, 1996; Irizarry et al., 2003; Gentleman et al., 2004). Genes that had the interquartile range (IQR) less than the median were detected as lowly varying, and were removed from further analysis of differential expression. To estimate differential expression among the remaining genes, a linear model was used (Smyth, 2004). The resulting *p* values of the t-tests were corrected for multiple hypothesis testing and reported as *q* values (Storey and Tibshirani, 2003). A threshold of *q* value < 0.01 was set to select significantly differentially expressed genes in response to drought.

### Functional and regulatory annotations of clusters

GO BP process enrichment analysis of clusters was performed using the cumulative hypergeometric test (as described above). For the webserver, GO BP genesets were downloaded from the PlantGSEA website (Yi et al., 2013). Terms that annotated more than 1500 genes and less than 10 genes were removed to gain resolution in the biological processes presented by enrichment analysis. CRE enriched in the clusters were identified using FIRE (Elemento et al., 2007). FIRE uses mutual information (MI) to find an association between the expression profile and the motif profile of genes, and uses randomization tests to score for statistical significance. Identified motifs present in more than 50% of all the genes in a given cluster were considered.

Perl scripts were used to parse all the data. Plots were generated using R (Ihaka and Gentleman, 1996) and gene expression matrices were visualized using MeV (Saeed et al., 2006). The gene expression data reported here are available from the NCBI GEO database with the accession number GSE81253.

## Results

### The rice environment coexpression network

To determine biologically meaningful stress transcriptional modules in rice, we have designed an extensive pipeline that uses data from publicly available gene expression profiles in parallel with our in-house generated datasets measuring drought response in three developmental stages (Fig. 1). We obtained 129 publicly available rice Affymetrix microarrays related to response of the rice plant to some environmental condition and worked with the raw data (Fig. 1, step 1a). The data was normalized and summarized into a gene expression matrix based on a custom probe-gene reannotation of the rice GeneChip. The reannotation increases the accuracy of the gene expression quantification process by assigning only specific probes to genes, and increases coverage of the array. The gene expression data was then converted into a matrix of 34,792 genes and 45 distinct conditions/groups and used to construct a coexpression network connecting pairs of genes that have a significantly high correlation between their expression profiles across the conditions (top 2.5% of all pairs of genes ordered in decreasing order of correlation; see Materials and Methods). This network, termed Rice Environment Coexpression Network or RECoN, contains 34,792 genes connected by ~18.5 million edges.

**Figure 1:**
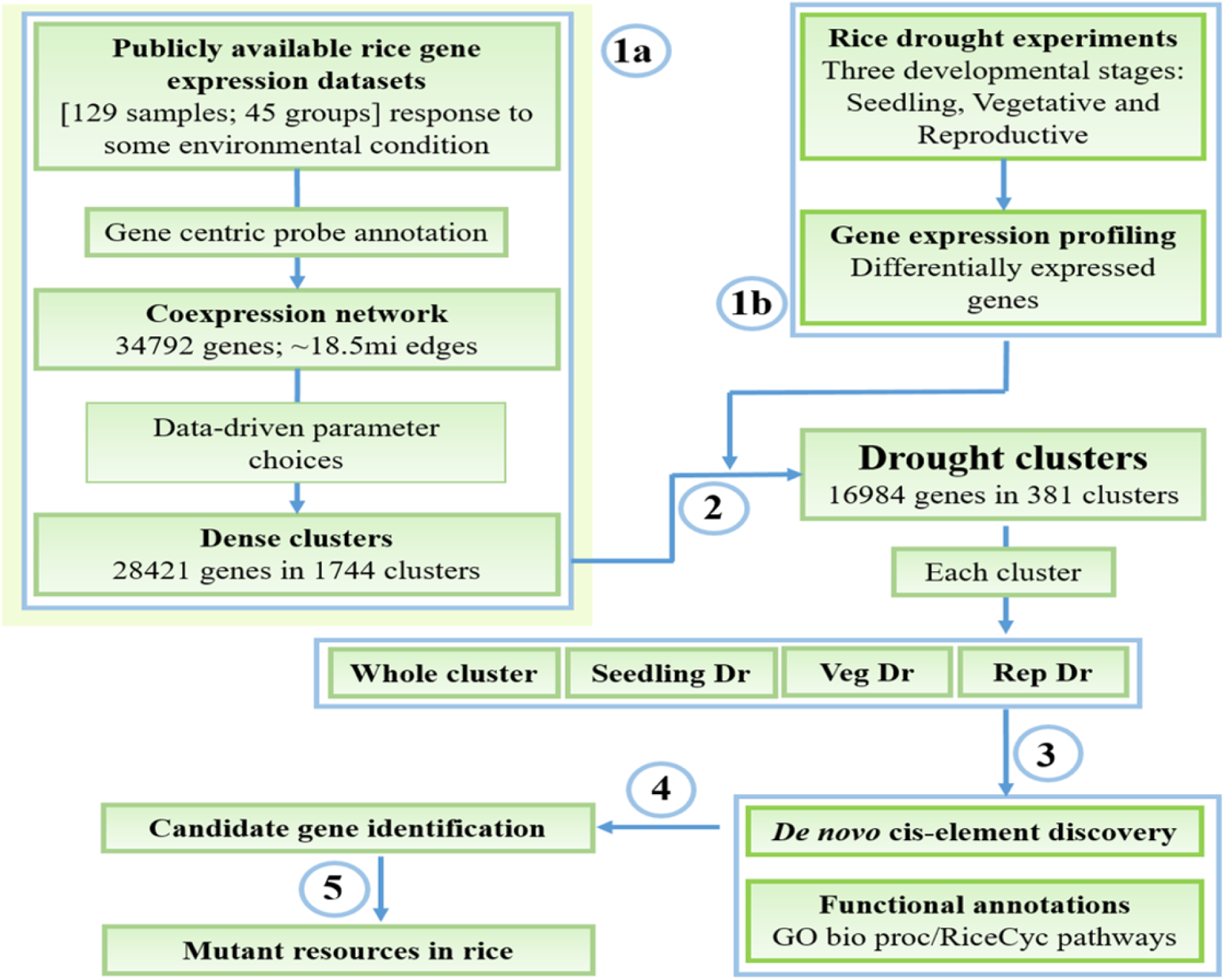
**Workflow for mining and characterization of drought transcriptional modules**. (**1a**) Reconstruction and clustering of the rice environmental coexpression network from publicly available gene expression datasets. (**1b**) Identification of drought-responsive genes in the three developmental stages. (**2**) Determination of ‘drought’ clusters based on the combination of results from the steps 1a and 1b, and extraction of whole cluster and specific drought gene sets (**3**) Functional enrichment analysis and cis-regulatory motif discovery. (**4**) Presentation of these data to the user where (s)he explores the results to identify candidates genes for functional validation. (**5**) Availability of mutants in genes of interest that can be used to study gene function

There are several clustering algorithms that work with weighted networks and find groups of densely connected nodes (Enright et al., 2002; Bader and Hogue, 2003). SPICi, a clustering tool was selected due to its ability to cluster large networks extremely fast (Jiang and Singh, 2010), and used to cluster our extremely large network. However, like every clustering algorithm, amongst a few, there is a single user defined parameter *T_d_* that determines the density of the resultant clusters and heavily influences the clustering process. To avoid an *ad hoc* or even a wrong choice of this parameter, we performed exhaustive data-driven tests on the network clustered using a range of *T_d_* values to identify the best parameter for the network at hand (Fig. 2).

**Figure 2:**
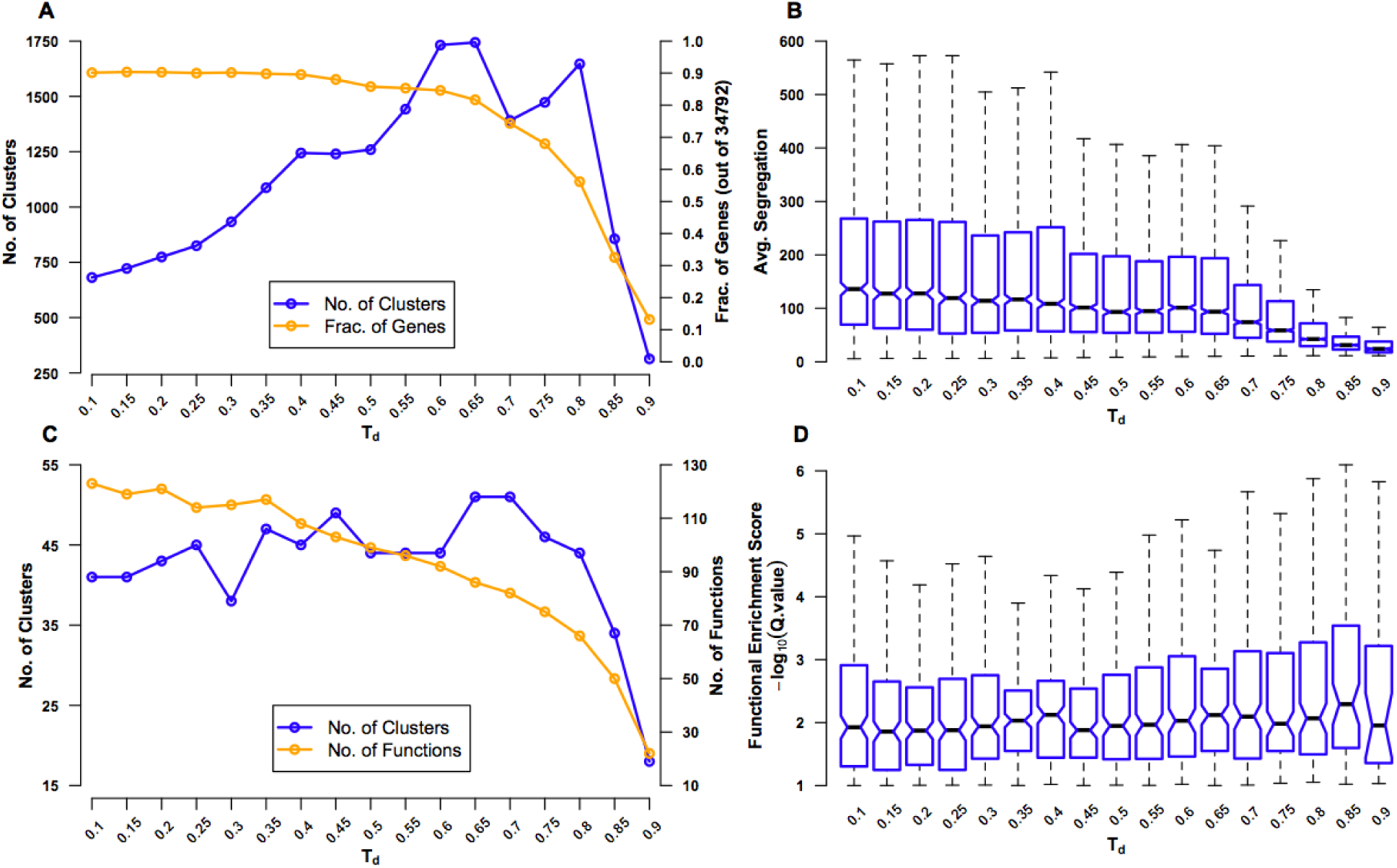
**Evaluation of coexpression network clustering**. The rice ‘environment’ coexpression network was clustered using SPICi, for a range of values – 0.1-0.9 – of the density parameter *T_d_* that determines how dense the final clusters are. The clusters obtained using each *T_d_* value were evaluated using several criteria: (**A**) Number of clusters that were formed (left y-axis) and the fraction of 34,792 genes in the original network present in one of the clusters (right y-axis) are plotted. These numbers were calculating by considering only clusters containing 3 or more genes. As *T_d_* increases, more and more genes are left out of clusters. (**B**) Average segregation of a cluster is a measure of how well genes in that cluster interact with other genes belonging to the same cluster compared to interactions with genes belonging to other clusters. Hence, average segregation measures cluster modularity. The overall modularity at a given *T_d_* value is plotted a box plot, leaving out outlier values above the whiskers for clarity. The center of the box corresponds to the median (2^nd^ quartile; *Q_2_*) of the distribution of average segregation values of all the clusters, and the extremes of the box correspond to the 1^st^ (*Q_1_*) and 3^rd^ (*Q_3_*) quartiles. The whiskers denote *Q_2_* ± 1.5\**IQR*, where *IQR* is the interquartile range (*Q_3_-Q_1_*). The notches in each box extend to +/-1.58 *IQR*/√*n* (*n* being the sample size) (McGill et al, 1978). They are based on asymptotic normality of the median and roughly equal sample sizes for the medians being compared, and are said to be rather insensitive to the underlying distributions of the samples. The notches give roughly a 95% confidence interval for the difference in two medians. (**C**) The extent of overlap between clusters (defined based on a particular *T_d_* value) and GO BP gene sets (termed ‘functions’) is measured using the hypergeometric test. The number of clusters with significant overlap (FDR *q-value* <0.1) (left y-axis) and number of distinct functions significantly overlapping with the clusters (right y-axis) are plotted. (**D**) Functional enrichment of the clusters is quantified using –*log_10_*(*q-value*) and plotted using a box plot representing the distribution of the enrichment scores for all the clusters at a given *T_d_* value. Here again, outliers beyond the whiskers have been left out for clarity

First, for different values of *T_d_*, we tracked the number of clusters obtained and the fraction of genes in the original network that were in clusters of 3 or more genes (Fig. 2A). At small values of *T_d_*, there are very few clusters and only a few broken links. As *T_d_* increases, the number of clusters increases, but, however, very high *T_d_* will break the network so much that the clusters with 3 or more genes will again become rare. Similarly, as *T_d_* increases, the number of genes that are part of clusters will steadily decrease until a critical value beyond which a large portion of genes will get disconnected and fall out of good-sized clusters. By testing for the value of *T_d_* after which there is the first significant drop in the number of clusters and fraction of genes in clusters, we found that this is at *T_d_*=0.65. Second, we calculated a measure of modularity called average segregation that quantified how well genes within a cluster are connected to each other compared to their connection to all the genes in the network (Fig. 2B) (Yook et al., 2004). Since we are interested in finding coherent biological modules, finding a *T_d_* that preserves segregation is sought after. It was surprising that the network showed the highest values of segregation for the smallest values of *T_d_*, indicating that even the original network with ~18.5 million edges is highly modular. Therefore, in the context of this network, at least, it was only important to look out for partitioning the network as much as possible without a significant drop in the inherent modularity. The first significant drop in average segregation (measured more qualitatively than quantitatively using the notches in the box plots; see Fig. 2 legend) occurs when the *T_d_* value is increased from 0.65 to 0.70, suggesting that setting *T_d_*=0.65 ensures the maximum modularity-preserving partitioning of the network.

Third, as we are interested in the functional consistency of genes within a cluster in addition to topological cohesiveness, we characterized the functional enrichment of all the clusters for a given *T_d_* value using GO BP enrichment analysis (Fig. 2C). Since this approach will suffer from the very sparse functional annotation of rice genes, we used this analysis only as a rough guide. Following the number of clusters that were significantly enriched with at least one specific GO BP (‘function’), we observed that the maximum enrichment again occurs at *T_d_*=0.65 (slightly better than *T_d_*=0.70). However, contrary to what is expected, the number of distinct enriched functions dropped steadily with increasing *T_d_*. Finally, using data from the enrichment analysis, we plotted the distribution of enrichment scores of all the clusters for different *T_d_* values and found that *T_d_* values in the range of 0.65 to 0.80 were giving overall more significant overlap between clusters and functions (Fig. 2D). Therefore, based on all the four analyses, we decided on a *T_d_*=0.65 to be the best choice for clustering RECoN.

We subsequently clustered RECoN using SPICi with *T_d_*=0.65 to uncover 1744 dense clusters with 3 or more genes. 28,421 genes (~81.7% of all the genes in the original network) fell within one of the clusters. Clustering the conditions based on their expression profiles also yields an expected grouping, especially with the drought-, salt- and cold stress samples clustering together (Supplemental Fig. S1). We linked these clusters to BP categories from the GO ontology and to CREs identified using a *de novo* motif discovery pipeline (Harb et al., 2010) (see “Methods”). The clusters thus identified can be used in a geneset enrichment analysis framework of new stress transcriptomes of rice. To demonstrate this analytical pipeline, we generated rice drought transcriptomes at three developmental stages, and used RECoN to identify clusters that are significantly perturbed in at least one stage.

### Gene expression profiling of drought in rice

We profiled RNA samples from rice plants treated to drought at the seedling, vegetative and reproductive stages using the rice Affymetrix GeneChips (Fig. 1, step 1b). In addition, we measured phenotypic and physiological responses of the plants to drought stress (see Supplemental Note and Supplemental Table S2). Statistical analysis of differential expression showed that a large number of genes are perturbed, given a stringent *q*-value cut-off of <0.01 (Supplemental Table S3). The largest shift in expression compared to well-watered controls happened at the seedling stage with ~12,300 genes showing differential expression, compared to only ~2,500 genes in the reproductive stage and ~9000 genes at the vegetative stage. A comparison of the differentially expressed genes at the three developmental stages showed that ~33% of the genes were shared with the genes in the other stages in the case of both up- and down-regulated genes (Fig. 3A and 3B).

**Figure 3:**
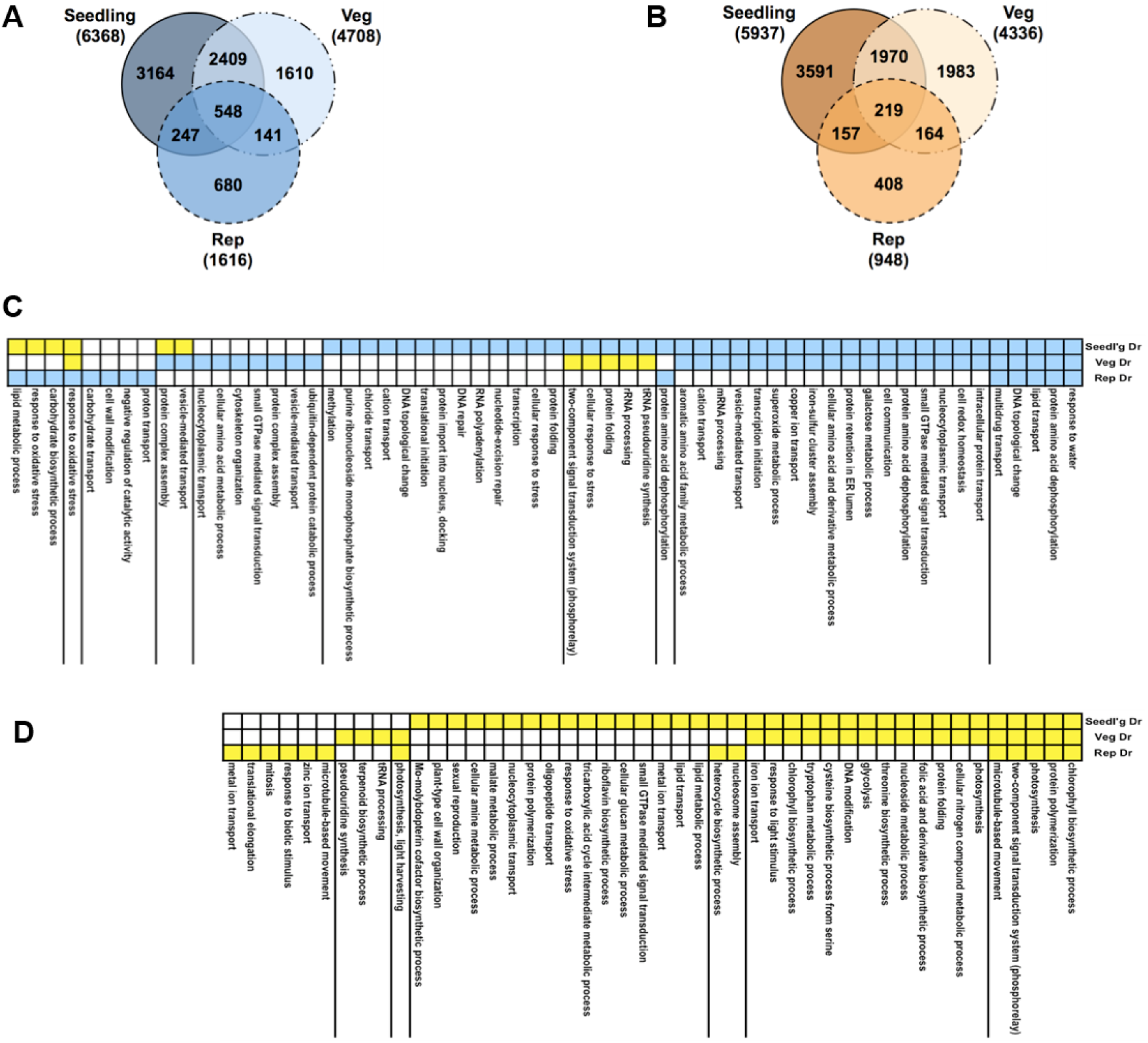
Gene expression profiles under drought. Venn diagrams comparing up-regulated (**A**) and down-regulated (**B**) genes in response to drought in three growth stages: seedling, vegetative and reproductive. Total numbers of genes for all gene sets are indicated in brackets. Functions, processes and pathways common and specific to various drought stress treatments and time-points. These are defined broadly based on Gene Ontology (GO) biological process (BP) annotations of rice genes. First, the total of all drought-regulated genes from all stages were pooled together and were then partitioned based on the combination of their regulation in the three stages (e.g. up-up-up, or down-up-down). Then, GO BP terms of interest (rows) were identified by analysis of enrichment of the set of genes annotated with a given GO BP term in each regulation-combination defined by the yellow-blue color-coding along the rows where blue means up-regulation and yellow means down-regulation. Statistical significance of enrichment was calculated using the hypergeometric test and terms with *q-value* <0.1 in at least one of the treatments were retained. (**C**) GO BP terms enriched in gene sets up-regulated in at least one stage. (**D**) GO BP terms enriched in gene sets only down-regulated in one or more stage.

To see the level of functional enrichment using genesets from the Gene Ontology (GO), we took a union of all the drought-regulated genes and split them into sets of genes that show identical pattern of regulation across the stages. We then determined the processes defined by GO biological process (BP) annotations that were enriched in each of these gene sets (Fig. 3C and 3D). As expected, the most significant GO term among the set of genes up-regulated in all stages was ‘response to water’. Similarly, different combinations of genes involved in protein dephosphorylation and small GTPase-mediated signaling are up-regulated in all stages. Among the genes down-regulated in all stages, photosynthesis and related processes are clearly enriched. Genes involved in translation are induced and repressed in the seedling and reproductive stages, respectively. Cell wall modification genes that are usually repressed by drought (Moore et al., 2008) are also repressed at the seedling stage but specifically up-regulated in the reproductive stage. Comparisons of the GO BP category revealed the most obvious differences between up- and down-regulated genes (Supplemental Table S4). In this category, response to water (GO:0009415), lipid transport (GO:0006869), cellular response to stress (GO:0033554), transcription (GO:0006350), response to oxidative stress and carbohydrate biosynthetic process (GO:0016051) were found at higher proportions in up- than in down-regulated genes. In contrast, the processes photosynthesis (GO: 0015979), chlorophyll biosynthesis (GO:0015995) and glycolysis (GO:0006096) were specifically represented in the down-regulated set of genes.

Although this analysis gave us a few insights into drought-regulated gene expression, apart from the *bona fide* stress response themes, it is hard to pinpoint biological functions that are specifically affected in the different stages. The most important reason for this shortfall is the fact that rice genes are extremely poorly characterized and very few genes have been annotated well. This scenario becomes evident when we look at the small number of genes common between any GO term and the set of drought genes. Therefore, we need to pursue other approaches that will give us a better picture of the underlying changes during drought.

### Identification of drought-related clusters from RECoN

The next operation was to interface the information gained from drought expression profiling to identify drought related modules from the coexpression data (Fig. 1, step 2). Each of the 1744 clusters from RECoN were tested for enrichment of drought-responsive (up- or down-regulated) genes from any one of the stages (seedling, vegetative or reproductive) (Fig. 4). Drought clusters provide a handle on putative functional interactions between genes transcriptionally regulated by drought that were otherwise unassociated parts lists. This makes gene-by-gene interpretation a much easier and constructive process. Moreover, we reasoned that since a cluster is a coherent group of genes, all the genes in a ‘drought’ cluster might have a role in mediating drought-response, not necessarily by responding to drought through gene expression changes. This is possible by either being ubiquitously present as support machinery (between well-watered and drought conditions) or being conditionally active under drought due to non-transcriptional modes of regulation including post-translational modification. These clusters, hence, provide a means for functionally associating post-transcriptionally modified regulatory/signaling genes to transcriptionally regulated genes.

**Figure 4:**
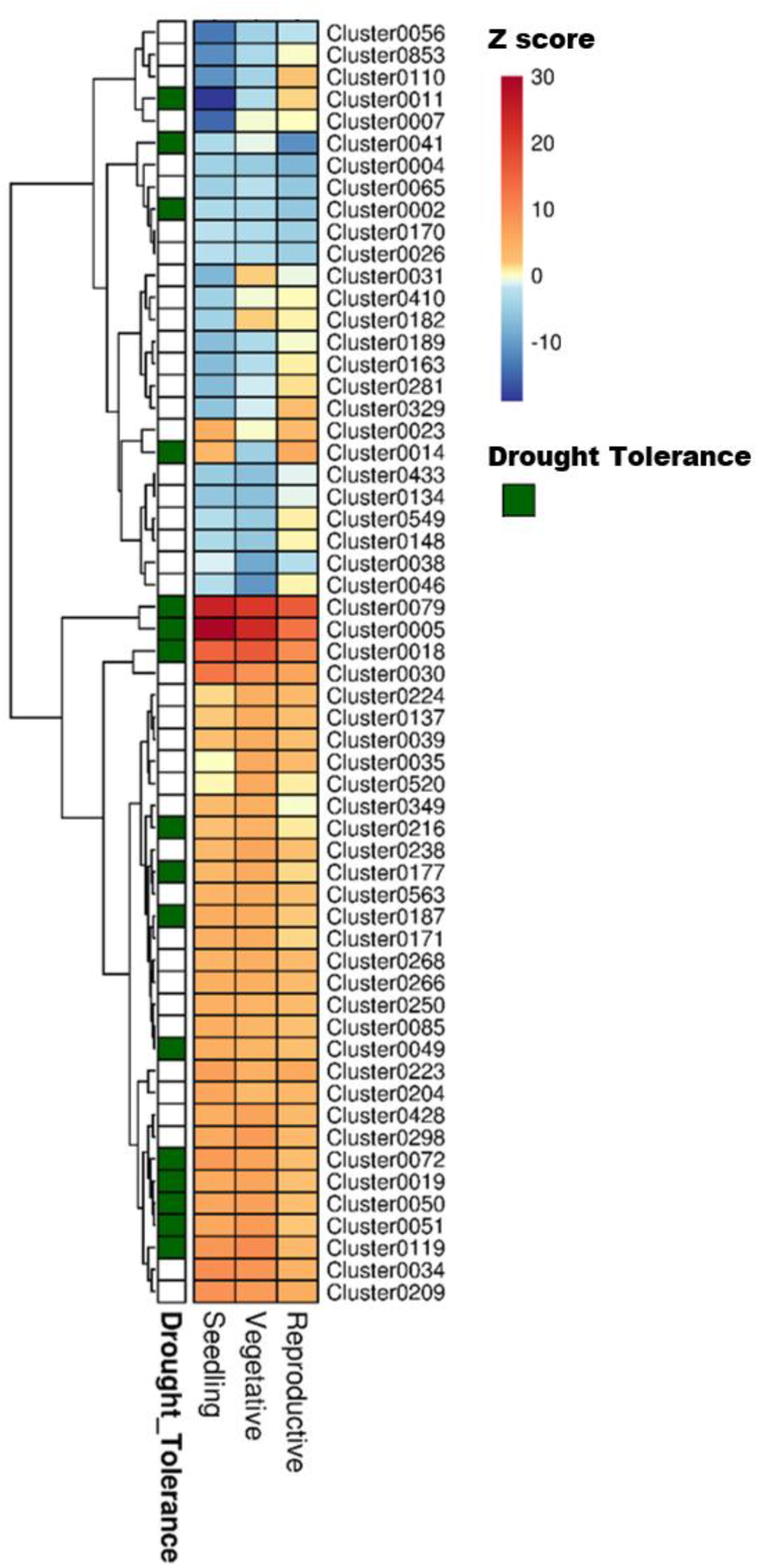
Heat map showing the different clusters enriched in drought responsive genes in three developmental stages of rice. The fold change values obtained from the differential expression tests was used as a parameter for the parametric analysis of geneset enrichment algorithm. The heat map shows Z scores obtained from the enrichment analysis, color coded with a red and blue gradient for positive and negative enrichment, respectively. The green grids along the rows indicate the ‘drought tolerance clusters’, identified by mapping known drought tolerance genes to clusters. A *q* value cut-off of 0.001 was set as a threshold to select the clusters.

### Examples of drought transcriptional modules

We present here some drought transcriptional modules as examples to showcase the usefulness of this approach in understanding developmental stage-specific drought response. All the genes in drought clusters, their relative expression across the stages and their ‘drought’ cluster membership are provided in Supplemental Table S5.

Cluster0013 contains 294 genes enriched with genes up-regulated in the seedling stage and down-regulated in the reproductive stage. Genes in this cluster are involved in ribosome biogenesis and mitochondrial protein localization (which concerns transporting of mitochondrial oxidative phosphorylation proteins to the mitochondrion), and contain the GCC-core, Telo-box and the Site II motifs in their upstream sequences. This combination of biological processes and CREs represents a well-known regulatory program: the site II motifs are recognized by TFs of the TCP family and have been confirmed to be important in the regulation of ribosome protein (RP) genes in combination with the telo-box motif (Tremousaygue et al., 2003). These motifs are co-located in the promoters of about 70% of 216 ribosomal protein genes in Arabidopsis. In addition, there is evidence that the site II motifs also possibly coordinate the expression of nuclear genes encoding components of the mitochondrial oxidative phosphorylation machinery in both Arabidopsis and rice (Welchen and Gonzalez, 2006). Therefore, this program involving site II and telo-box motifs could mediate the down-regulation of major processes that affect protein production under drought stress in the reproductive tissue. The GCC-core motif is known to be bound by AP2-ERF TFs (Ohme-Takagi and Shinshi, 1995), which are involved in gene regulation under a variety of abiotic stresses conserved between Arabidopsis and rice (Nakashima et al., 2009).

Cluster0010 contains 635 genes including genes involved in lignin biosynthetic process, amino acid transport, systemic acquired resistance, glycolysis, pentose-phosphate shunt and two-component signal transduction system (phosphorelay). Genes in this cluster are down-regulated in the seedling and vegetative stages, but up-regulated in reproductive stage. Of particular interest in this cluster is the *OsVIN1* gene (LOC_Os04g45290) coding for a vacuolar invertase gene. OsVIN1 has high fructan exohydrolase activity and is known to play an important role in carbon allocation to developing organs like the reproductive tissue. The expression of *OsVIN1* is not induced by our drought treatment, and this is in agreement with previous observation that *OsVIN1* is expressed in flag leaves, panicles (the reproductive tissue) and anthers in an essentially drought-insensitive manner (Ji et al., 2005; Parent et al., 2009). It is therefore a case where a gene involved in mediating a process (resource allocation) relevant to drought is not transcriptionally affected, but is associated with other drought-regulated genes in clusters defined by us. Another important gene observed in this cluster is *SNAC3* (LOC_Os01g09550) that has been previously shown to confer tolerance to multiple stresses like salinity, drought and oxidative stresses (Fang et al., 2015). *OsCPK9* (LOC_Os03g48270), a calcium dependent protein kinase is another key member in this cluster that has previously been shown to impart drought tolerance in transgenic rice plants by enhancing stomatal closure and stomatal adjustment. (Wei et al., 2014). Taken together these results suggest that the genes in this cluster contribute to drought tolerance by regulating osmotic adjustment and ROS scavenging processes and can also be putative candidates for increasing yield under drought.

The 193 genes in Cluster0041 are enriched primarily in almost all processes involved in cell cycle, a process integral to panicle development and elongation, and these genes are specifically down-regulated by drought at the reproductive stage (the most drought sensitive stage of rice). Upstream regions of these genes contain the *SEF3* binding site/ACII element, *MYB* recognition site found in *rd22* and other genes, and E2F consensus, potential binding sites of TFs that have been implicated to be important in regulating cell cycle in the reproductive tissue of Arabidopsis (Hennig et al., 2004).

The other aspect of using this approach is in discovery of drought tolerance genes. A variety of gene families with regulatory function have been shown to have a role in drought tolerance by overexpression/knockout experiments, and that regulate a battery of downstream genes (Umezawa et al., 2006). Therefore, to evaluate this aspect, we first catalogued a number of genes that confer drought tolerance in rice on overexpression or knockout, and then mapped them to RECoN clusters (Fig. 5). The primary observation is that almost all the drought tolerance genes were part of drought clusters. However, this observation could be trivial if all those genes are indeed regulated by drought in the first place. Out of the 54 genes presented here, 45 are indeed regulated by drought in stage specific or independent manner while 9 of these are not drought-regulated, but are associated with a drought module. Therefore, we reaffirm that the approach lends itself to identification of genes that are not necessarily transcriptionally perturbed by drought, if at all regulated by it. Some examples for the drought-tolerance clusters follow.

**Figure 5:**
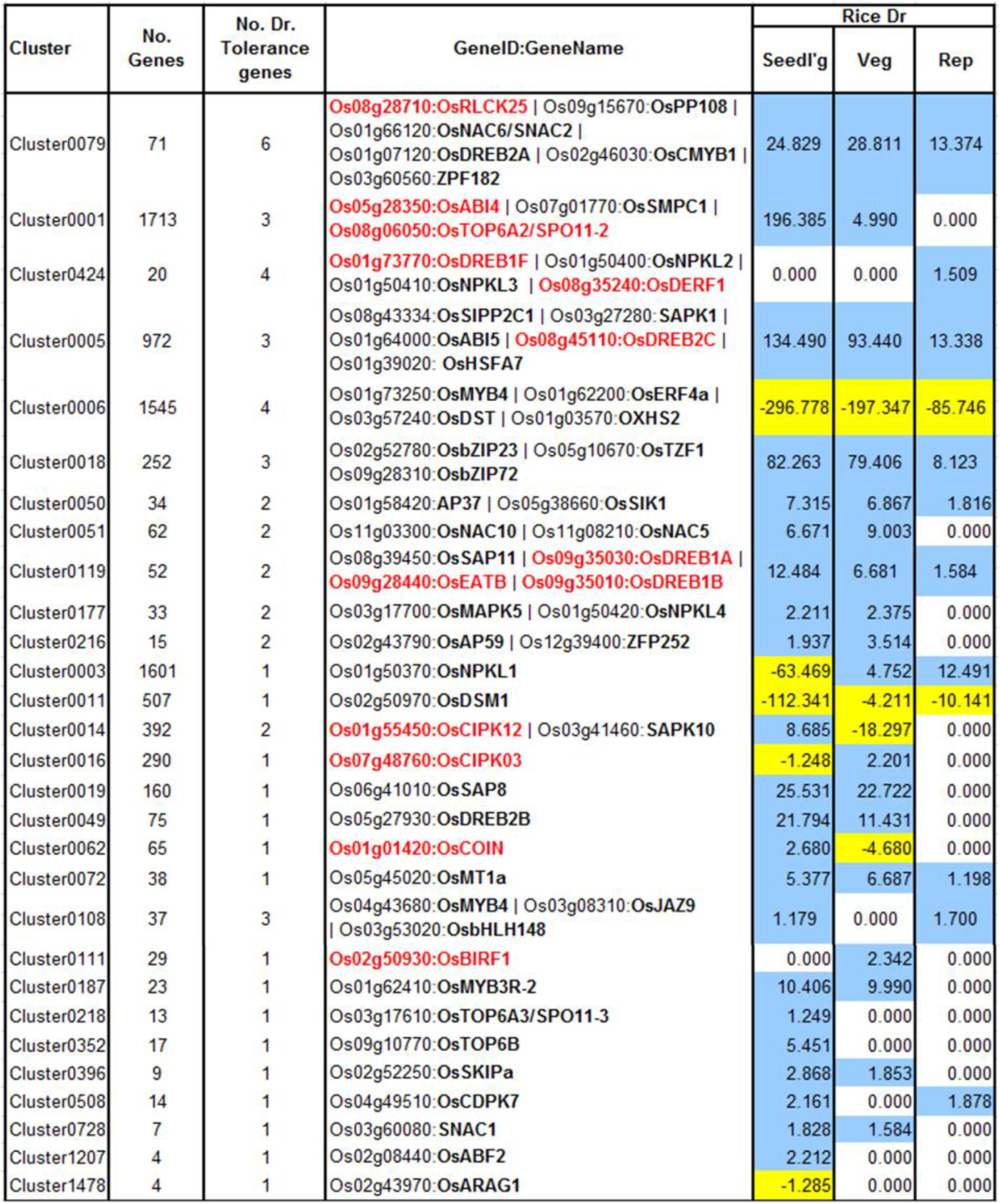
Drought clusters containing known drought tolerance genes. Genes in black are regulated by drought at one or more of the growth stages while genes in red are not drought-regulated. The values in the color-coded columns correspond to the level of significance (measured as score equal to the –*log10[q-value]*) of drought-regulated genes. For convenience the scores themselves are signed and colored based on the direction of their regulation (+/blue - up-regulation; –/yellow - down). Since only enrichments with *q-value*<0.1 were considered, all the other values were set to 1 (because of which, their negative logarithms are 0s).

Cluster0079 contains 71 genes. The genes in this cluster include a receptor-like cytoplasmic kinase *OsRLCK253* (LOC_Os08g28710) and a phosphatase *OsPP108* (LOC_Os09g15670), both of which have been shown to improve drought tolerance in transgenic *Arabidopsis* plants (Giri et al., 2011; Singh et al., 2015). Along with the aforementioned genes, this cluster also comprises of other known drought tolerance genes comprising of dehydrins like *OsLea3-1* (LOC_Os05g46480) (Brohee and van Helden, 2006), enzymes like *OsUGE-1* (LOC_Os05g51670) (Nardini et al., 2011) and TFs like *OsDREB2A* (LOC_Os01g07120) (Skirycz and Inze, 2010), *SNAC2/OsNAC6* (LOC_Os01g66120) (Bergmann et al., 2004), *CMYB1*(LOC_Os02g46030) (Kitano, 2002) and *ZFP182* (LOC_Os03g60560) (Fig. 6A). Most genes in the cluster are up-regulated by drought in all three developmental stages, which appears to indicate the diverse roles that these proteins play including detoxification, osmotic adjustment, and signaling pathways. Since our data revealed many putative stress inducible genes, a few of these genes are likely to have a dual role as developmentally regulated and stress responsive. Nevertheless, the functional role of these genes needs to be characterized to further enhance our understanding of the mechanisms that impart drought/abiotic stress tolerance to rice.

**Figure 6:**
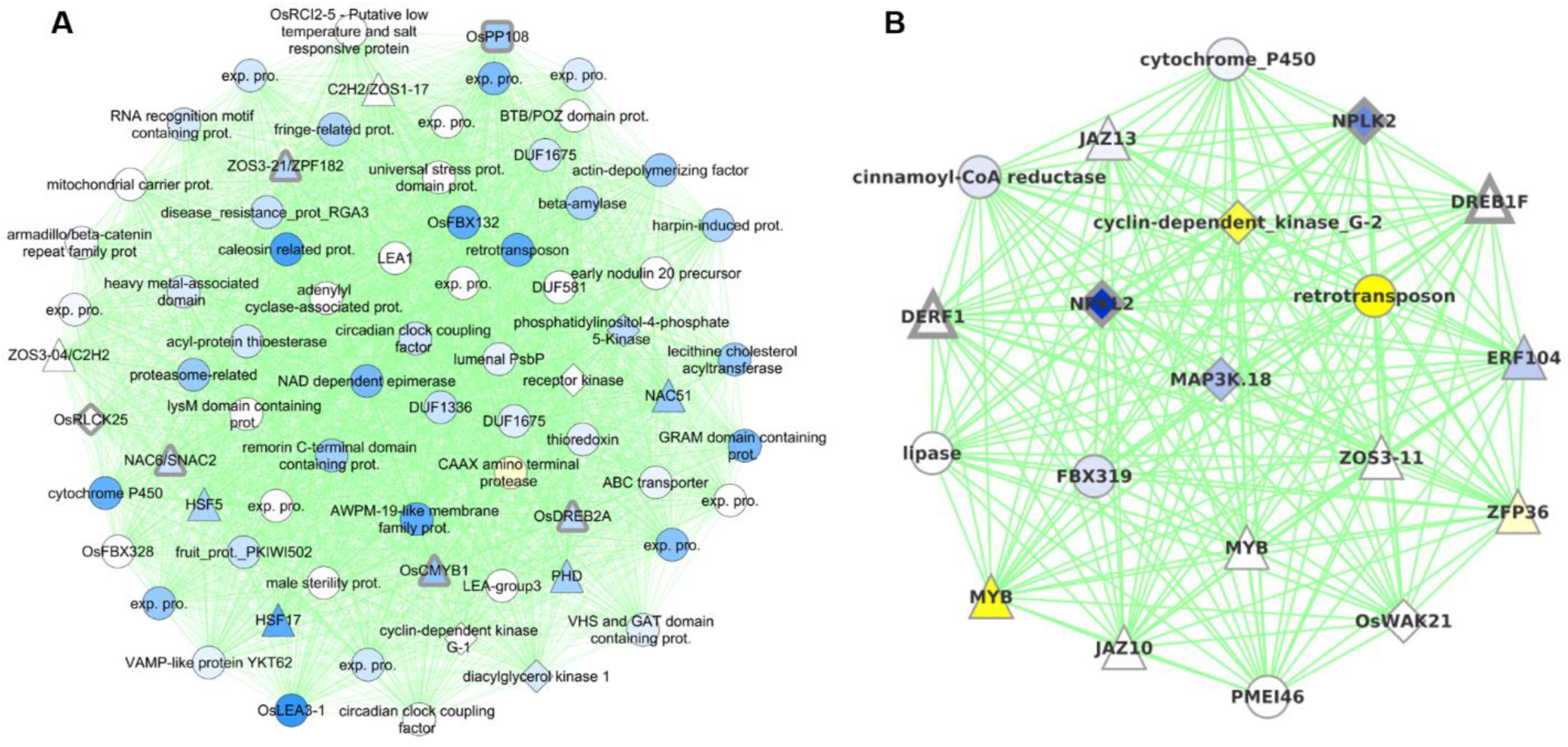
Graphical visualization of A) 71 genes in Cluster0079 that contains six drought tolerance genes (with thick grey borders) and B) 20 genes in Cluster0424 that contains four drought tolerance genes. All the coexpression edges are colored green. Node shapes correspond to type of gene: triangles are TFs, diamonds are protein kinases, rounded squares are protein phosphatases and circles are other genes. Node color corresponds to the level of differential expression under drought in the vegetative stage for Cluster0079 and reproductive stage for Cluster0424 (where the clusters have maximum enrichment): blue for up-regulation and yellow for down-regulation. Uncharacterized gene are labeled ‘exp. pro.’ (for ‘expressed protein’).

Cluster0424 contains 20 genes enriched specifically with reproductive drought and these genes too contain an ABRE-like motif – HACGYGTNS – in their upstream sequence. The drought tolerance genes part of this cluster are *OsDREB1F* (LOC_Os01g73770) (Barrero et al., 2007; Choi et al., 2007), *OsNPKL2* (LOC_Os01g50400) and *OsNPKL3* (LOC_Os01g50410) (Fig. 6B). The tandem duplicate genes *NPKL2* and *NPKL3* are previously known to be strongly induced by drought at the reproductive stage (Ning et al., 2008). Highly induced expression of these genes under drought stress indicates that these two genes can be candidates for drought tolerance and increased yield under drought, for their potential role as a kinase as they are found to be located in the genomic region with three QTLs: RSN (relative number of spikelets per panicle under drought stress) and LDS (leaf drying score), which are mainly related to drought tolerance, and DIDRV (deep root rate in volume induced by drought conditions) (data not shown). Although expression of *OsDERF1* is induced by drought and phyothormone treatments, its overexpression in rice negatively regulates drought tolerance by repressing ethylene biosynthesis by binding to ERF repressors *OsERF3* and *OsAP2-39* (Wan et al., 2011). *OsDREB1F* was induced by abiotic stresses including osmotic stress (using PEG) as well as ABA treatment and overexpression in rice and *Arabidopsis* gave drought tolerance that can be mediated by ABA dependent pathway (Choi et al., 2007). However, our progressive drought treatment does not perturb this gene (at least not at the stringent level of significance chosen). Developmental stage-specific drought-regulation of *OsDREB1F* is not clear except that the gene by itself is expressed differently in different stages and tissues. We therefore implicate *OsDREB1F* as being important in progressive drought response at the reproductive stage.

Cluster0177 contains 33 genes involved in the regulation of innate immune/defense/stress response as well as response to jasmonic acid and salicylic acid. Drought regulated genes in this cluster are up-regulated specifically in the seedling and vegetative stages. This cluster again contains the drought-tolerance genes *OsMAPK5* (LOC_Os03g17700) and *OsNPKL4* (LOC_Os01g50420). *OsMAPK5* is known to be induced by drought, other abiotic stresses and ABA, as well as pathogen infection and that the overexpressiwon gives abiotic stress tolerance but disease susceptibility (Xiong and Yang, 2003). It is hence considered to be a key link in the cross talk between disease resistance and abiotic stress tolerance. We propose that other genes in this cluster are putative links of crosstalk between the stresses. Previous research has shown that *OsNPKL4* is very strongly induced at the seedling stage, but has a moderate to low level of induction at the anthesis stage (Ning et al., 2008), consistent with the drought-pattern of this cluster.

Cluster 0108 contains 37 genes with three drought tolerance genes. *OsMYB4* (LOC_Os04g43680), characterized as a universal stress response gene induced under a variety of biotic and abiotic stresses gives abiotic stress tolerance when overexpressed in apples (Yu et al., 2006; Narsai et al., 2013) by modulating osmolytic balance, *OsbHLH148* (LOC_Os03g53020) and its interacting partner *OsJAZ1* (LOC_Os03g08310) mediates drought response via the jasmonic acid pathway (Seo et al., 2011). Genes in this cluster also could potentially be involved in jasmonic acid mediated hormonal signal transduction.

### A web interface for further exploration of rice abiotic stress response

We have made RECoN available online at https://plantstress-pereira.uark.edu/RECoN/ (Fig. 7). The interface provides an enrichment tool (Kim and Volsky, 2005) that allows users to upload their own rice genome-wide transcriptional response and explore stress clusters significantly perturbed in their experiment (using the strategy outlined in Figure 1 and used in this study). Users can explore the pathways/processes and regulatory sequences linked to each cluster as well as the genes within that cluster reported along with their perturbation in the user data and our drought experiments, homologs in Arabidopsis, and available gene annotations from both the MSU release 7 (Kawahara et al., 2013) and RAP (Sakai et al., 2013) databases (Supplemental Fig. S2, S3 and S4). The webserver is Cytoscape-web (Lopes et al., 2010) enabled and allows graphical display of clusters with node attributes set to highlight changes in the user provided transcriptome and genes with regulatory roles (e.g. TFs and kinases). The platform also allows users to query a single gene as guide and retrieve a coexpression neighborhood to explore their functional context. Together, these functionalities in RECoN will enable biologists gain a network-based understanding of the stress response in rice and prioritize candidates for studying experimental phenotypes.

**Figure 7:**
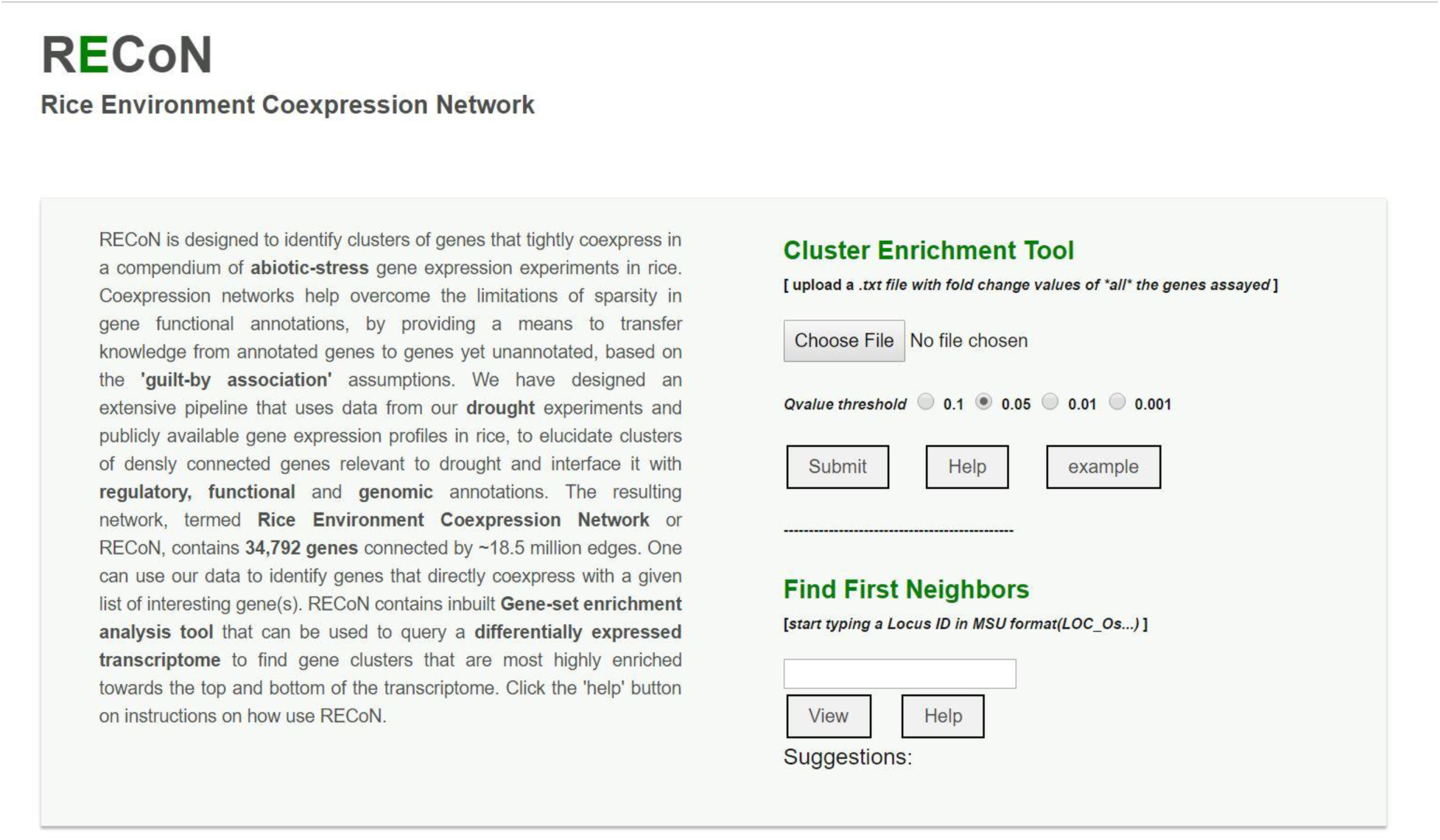
A screenshot of the RECoN webserver available at the link provided in the main text. The online platform allows two types of analyses. The user can upload a genome-wide differential expression profile using the ‘choose file’ option, which will be used by the cluster enrichment tool to identify clusters that are significantly perturbed in the uploaded transcriptome, within the selected *q* value threshold. The uploaded file should contain two columns (with headers) with MSU formatted rice gene locus IDs in the first column and their respective fold change values determined from the differential expression tests in the second column. The results will be displayed in a new page with enriched clusters listed and links to display each cluster using Cytoscape-web, as well as biological processes and *cis* regulatory elements enriched in the clusters (see supplemental figures S2, S3 and S4). In cases where a single gene is of interest (rather than a genome-wide analysis), its locus ID can be entered in the input box under the ‘Find First Neighbors’ section. This analysis will report the genes within one path length of the query gene and a default coexpression score of 0.80 (which can be changed from the results page, see supplemental figure S5).

## Discussion

Plant responses to environmental stress span across several layers of organization including signaling, transcription, and metabolism, making it vital to understand stress response at the systems-level. For less studied models like rice, the current scope for systems analysis is mostly restricted to transcriptional profiling under various conditions. Therefore, to make the best use of currently available data in rice, we have created a resource for exploration of transcriptional, developmental, functional, and regulatory aspects of abiotic-stress response in rice.

We sought to organize genes into coherent groups and work further from there. To this end, we designed and implemented a pipeline for automatic mining of condition-specific gene expression datasets intended for analysis of coexpression. At a practical level, accurate quantification of gene expression using technologies like Affymetrix GeneChips has been hard due to the problem of cross-hybridization. This has been noted to affect calculation of coexpression (Casneuf et al., 2007) and the proposed solution is a remapping of microarray probes to genes to ensure unique hybridization (Dai et al., 2005). We hence used a custom probe-gene mapping and used this reannotation to make reliable estimation of gene expression across 45 conditions. Then, a coexpression network was built (RECoN) and clustered to obtain tightly coexpressed groups of genes that revealed the modular organization of genes.

We demonstrated the use of RECoN by analyzing new stage-specific drought transcriptomes. In order to both understand drought response and discover novel drought tolerance genes, we combined drought-responsive genes from our experiments with the transcriptional modules to uncover drought clusters, where each cluster, by design, contains several genes in addition to genes transcriptionally regulated by drought. Drought modules thus present an opportunity to discover regulatory genes that do not change in gene expression but can affect the response mediated by that module. In this process, we are basically imputing uncharacterized genes within a cluster with the function/role of characterized genes (even at the level of transcriptional response). In species with very little annotation, such as rice, cluster-level function prediction has been shown to be useful (Song and Singh, 2009). We have validated this approach by inspecting the cluster membership of known drought tolerance genes that are not drought responsive but are associated with a cluster that is enriched in genes following a drought expression pattern expected from what is known about the tolerance gene.

With the enormous amount of data generated in this work that can be used for inference of gene function and pathway analysis, all these results are summarized and presented in a flexible visual interface for dynamic exploration. This online platform is – to the best of our knowledge – the first of a kind that allows users to upload their own transcriptomic data (e.g. output of an RNA-seq assay) and find clusters that are significantly enriched. The clusters are linked to GO BPs and CRE, eliminating the need for a traditional GO BP enrichment analysis. The approach presented here is widely applicable: genome-wide transcriptional modules recovered here on the basis of gene expression under different environmental conditions can be similarly extended to study other abiotic stresses including salt and cold to find common stress-specific modules. This approach lends itself to identification of stress-related genes that are usually hidden in a typical transcriptome assay.

## Acknowledgements

We thank members of the Pereira lab- Supratim Basu, Venkategowda Ramegowda, Ritu Mihani and Julie Thomas for testing and validating the network clusters for gene function analysis, some of which are in publications. We also thank Pawel Wolinski from the Arkansas High Performance Computing Center for his contribution to creation of webserver.

